# Development of a Novel Index to Characterise Arterial Dynamics Using Ultrasound Imaging

**DOI:** 10.1101/2020.10.31.363291

**Authors:** Joel Ward, Xinghao Cheng, Yingyi Xiao, Pierfrancesco Lapolla, Anirudh Chandrashekar, Ashok Handa, Robin A Cleveland, Regent Lee

## Abstract

Abdominal aortic aneurysms (AAA) are associated with systemic inflammation and endothelial dysfunction. We previously reported flow mediated dilatation (FMD) of the brachial artery as a predictor of AAA growth. We hence hypothesised that other physical characteristics of the brachial artery correlate with AAA growth. Using a prospectively cohort of AAA patients, we devised a ‘brachial artery relaxation index’ (BARI) and examined its role as a biomarker for AAA growth. However, no correlation between BARI and future aneurysm growth was observed (p=0.5). Therefore, our investigations did not substantiate the hypothesis that other physical characteristics of the brachial artery predicts AAA growth.

## Introduction

Abdominal aortic aneurysms (AAA) are permanent and irreversible localised dilatations of the aorta involving all three layers of the vascular wall. The incidence of AAAs increase with age demonstrated through ultrasonography based screening studies diagnosing AAA in 1-2% of all 65−year-old men and in 0.5% of 70−year-old women ^1, 2^. AAAs are usually asymptomatic so are most frequently detected either as part of screening programmes or incidentally on presentation to healthcare for an alternate pathology. AAAs have a tendency to expand, and as the size increases, so does the risk of rupture. When a AAA ruptures catastrophic intra-abdominal haemorrhage ensues which, if left untreated, results in mortality.

International guidelines recommend the A-P diameter threshold of 55mm for elective surgical repair in men, at which point the yearly risk of rupture outweighs the risks associated with an operation ^3^. Accordingly, patients with AAAs less than 55mm are typically kept under surveillance by regular ultrasound until the AAA expands above the threshold. Longitudinal studies have demonstrated that AAAs grow at different rates in different individuals and possess a variety of different physiological characteristics, the subtleties of which are not incorporated in current decision-making guidance ^4, 5^. Although 70% of patients eventually meet the size threshold for an operation, they age and become more co-morbid during the observational period, raising the risks associated with intervention ^4, 5^. Robust biomarkers of future AAA progression can improve clinical management by aiding identification of aneurysms which will progress to the necessitating intervention without requiring delay caused by a period of observation.

AAAs are associated with features of systemic inflammation and endothelial dysfunction ^6-8^. Endothelial dependent vasomotion has been widely used as a surrogate marker of endothelial function. Flow mediated dilatation (FMD) is a repeatable, non-invasive physiological assessment for the quantification of systemic endothelial function which has been shown to be inversely correlated with future AAA progression in humans ^9^. FMD deteriorates during the natural history of AAA, and is improved by surgery in a fashion predictable with modest accuracy ^9^.

We sought to improve the predictive power by incorporating further biomechanical data extractable from the ultrasound data loops already recorded for FMD to find another novel biomarker during the natural history of individuals with AAAs. We hypothesise that intrinsic physical viscoelastic characteristics of the brachial artery wall correlate with AAA growth.

## Material and Methods

Details regarding the OxAAA study cohort and recruitment process have been published ^9^. This prospective study received full ethics approval (Ethics Ref: 13/SC/0250) and recruited patients who received AAA monitoring in the National Health Service setting. Healthy volunteers without a diagnosis of AAA were also recruited as a comparison. Every participant provided written informed consent to participate in the study.

The AAA size was defined by the maximal anteroposterior diameter (outer-to-outer). We defined further subgroups according to the AP diameter (APD) into small (30-39mm), moderate (40- 55mm), and large (>55mm) AAAs. The annual growth rate of AAA during surveillance was calculated by: (∆APD/APD at baseline) / (number-of-days-lapsed/365days).

The research appointments for each patient took place on the same visit as the NHS AAA surveillance scan, during daytime hours. B mode ultrasound imaging of the brachial artery was then performed using the CX50 ultrasound machine (Phillips, Amsterdam, Netherlands) with a L12-3 probe which was mounted using a bespoke holder (**Figure 1**) to record brachial artery motion during four cardiac cycles. All data were anonymised before use in the work reported in this article.

**Figure 1:**
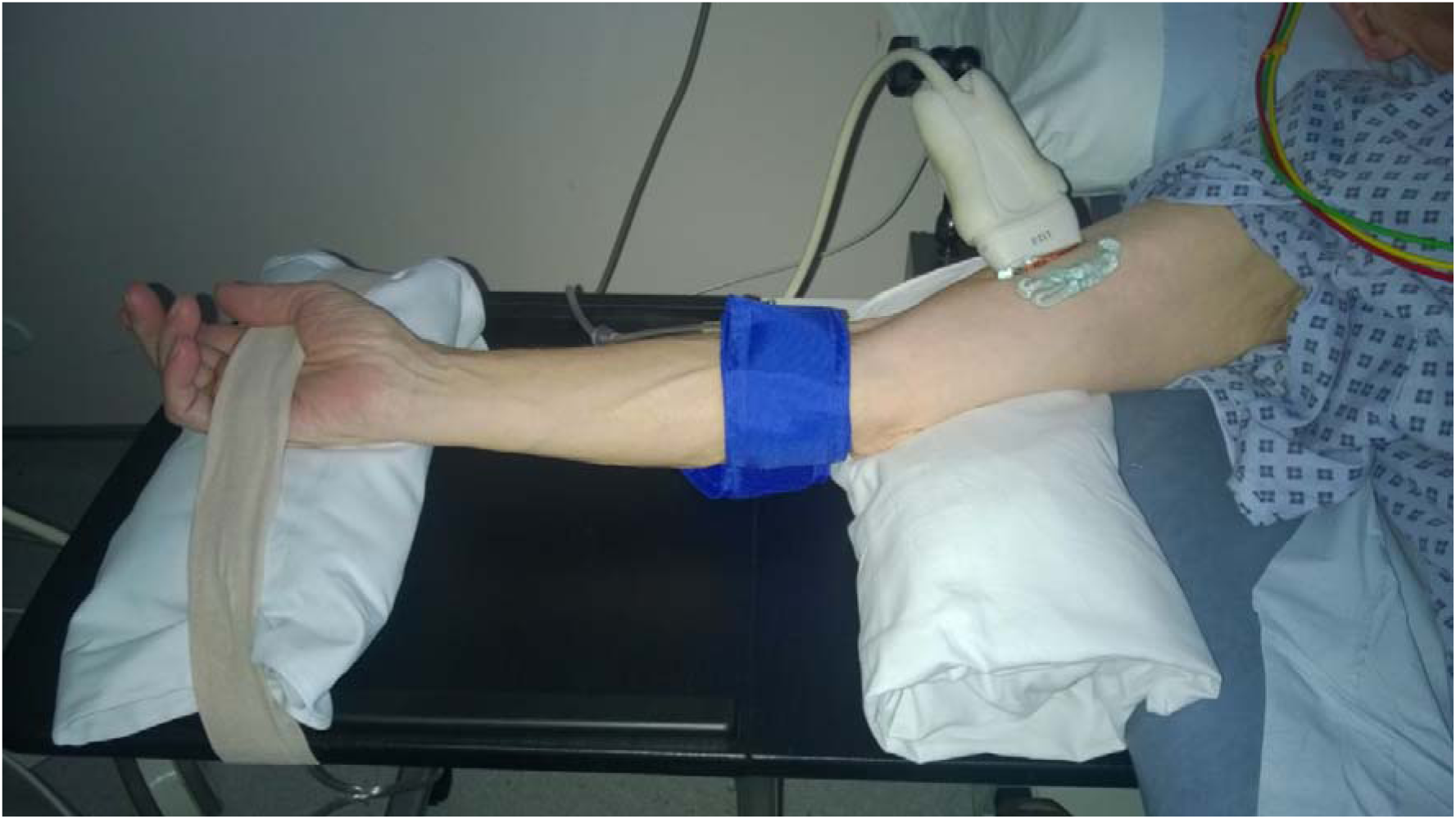
Set up to record brachial artery ultrasound video loop.

### Image Analysis

Ultrasound recordings were saved in DICOM format and imported into MATLAB® (The MathWorks Inc. Natick, Massachusetts, USA) for analysis. Brachial artery wall positions were detected using a modified version of a level set evolution (LSE) approach traditionally used in MRI information processing ^10^. This LSE method exploits the difference in pixel intensity in the grey scale ultrasound to locate the contours. A user-defined rectangle was manually placed within the lumen position in the first frame of each ultrasound recording (**Figure 2**, panel A). The algorithm then iterated outwards towards the arterial wall detecting differences based on the brightness intensity contrast. The algorithm was designed to find the contours of the vessel outside the initial user- defined rectangle. Care was taken such that the rectangular region of interest is not close to lumen edges to prevent overlap with the arterial wall during cardiac cycles.

**Figure 2.**
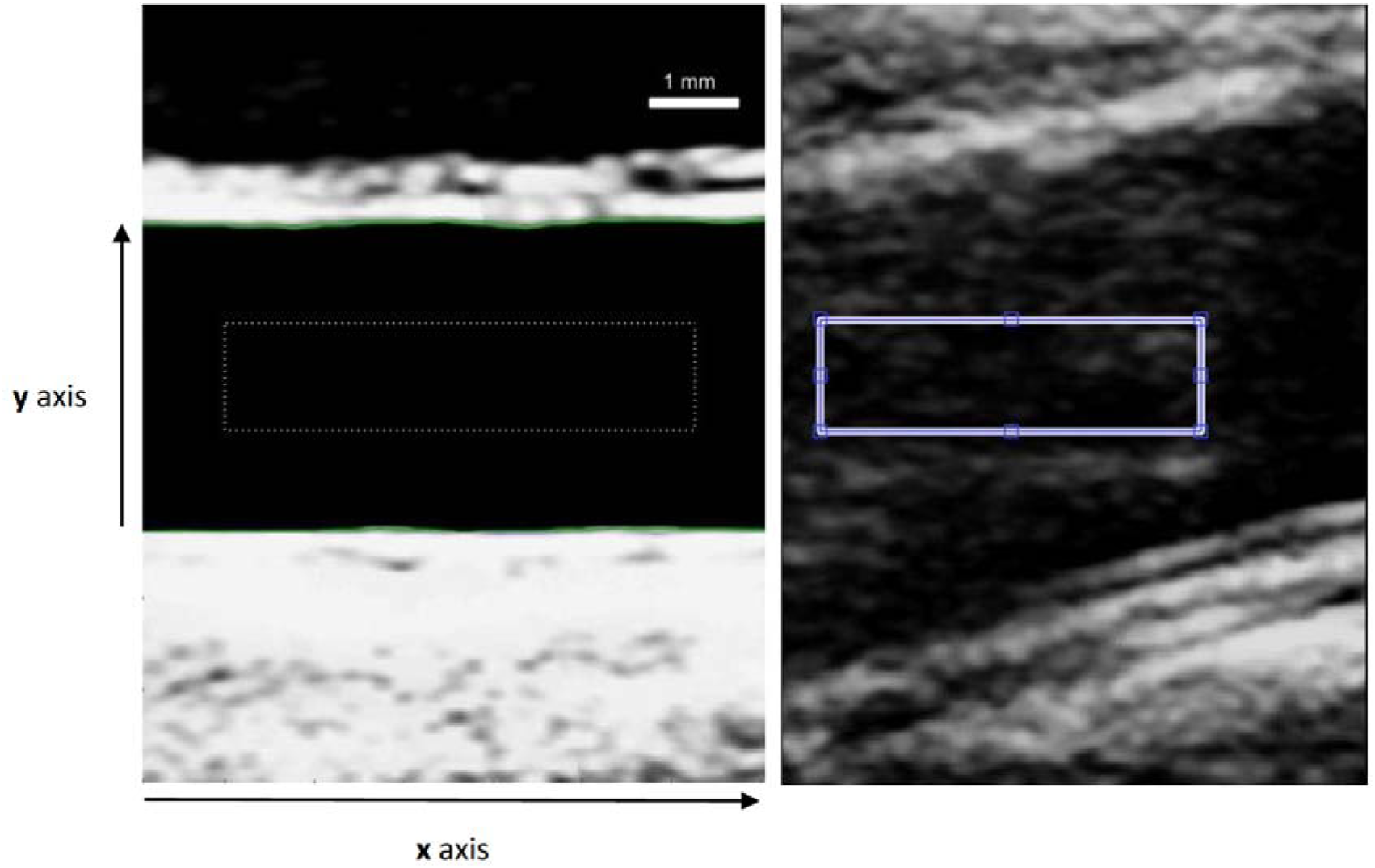
Image of a frame extracted from the ultrasound video of a brachial artery. The dotted rectangle is a user specified initialisation for the level set algorithm. **(A)** The green lines show the final estimate of the anterior and posterior vessel wall from the level set algorithm **(B)** Additional slice thickness artefact densities present in vessel lumen.

At each horizontal location in the image (x), the top of the lumen yt(x) and the bottom of the lumen *Yb*(x) were determined. This was repeated for each image in the cine loop which normally contained four cardiac cycles. **Figure 3** shows an example surface plot of *y*_*B*_ *(xH)ad y*_*t*_ *(x,t)*where four cycles can be seen and the response is reasonably uniform along the length of the artery that was imaged. The diameter D as a function of x was determined by:

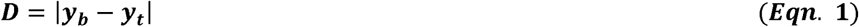

At each location x, the diameter D(x,t) was manually using the peaks and troughs so that systole and diastole could be identified, see **Figure 4**. The diameter of the vessel dilatation during systole was fit to a single exponential rise time.

**Figure 3.**
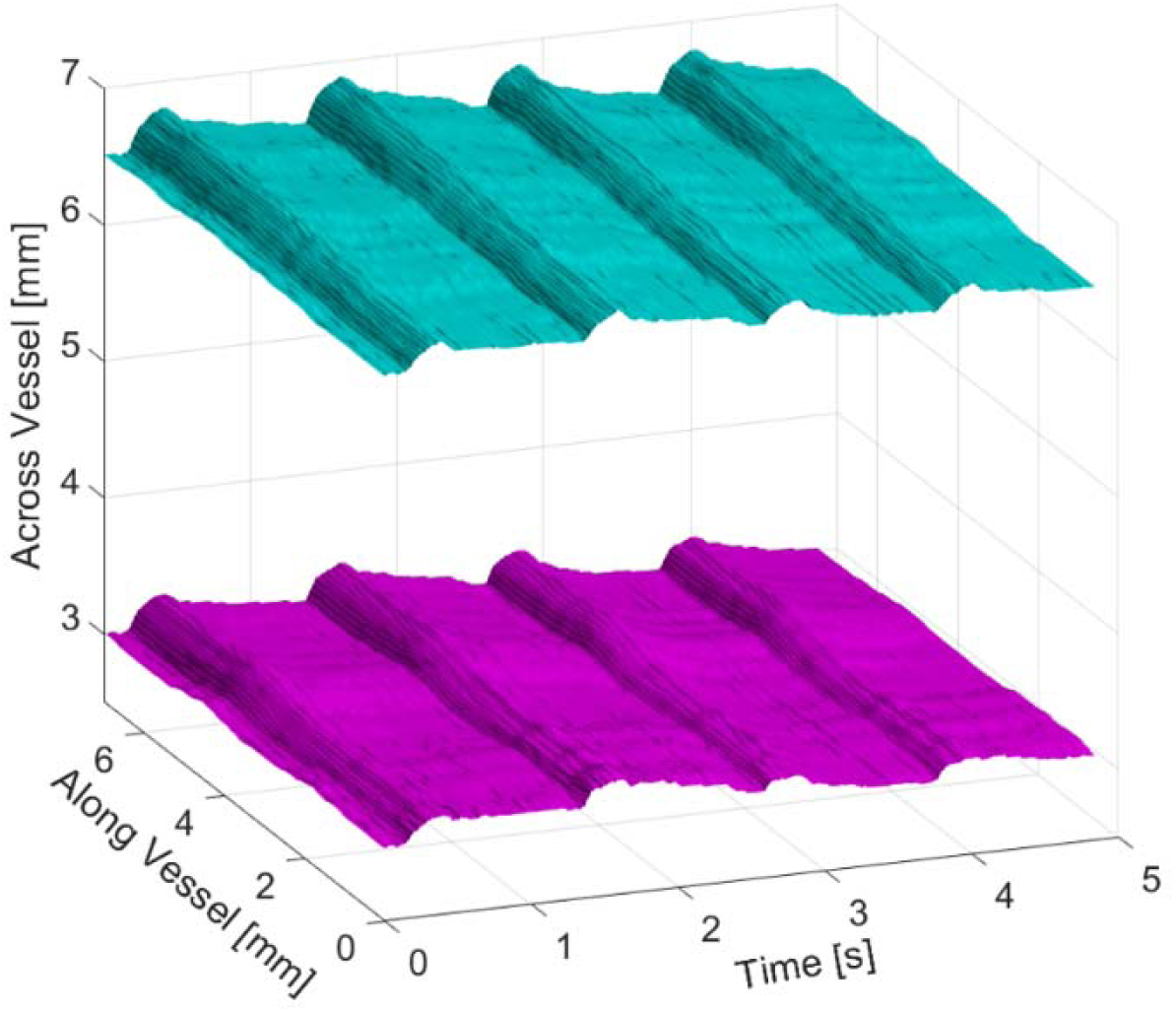
Response of brachial vessel wall in time and space over four cardiac cycles. The contour acquisition is conducted by the segmentation algorithm throughout the period of the ultrasound video clip. The expansion of the vessel during systole can be seen at about 0.5 s, 1.5 s, 3.0 s and 4.0 s. Cyan contours represent the posterior vessel wall. Magenta contours represent the anterior vessel wall.

**Figure 4.**
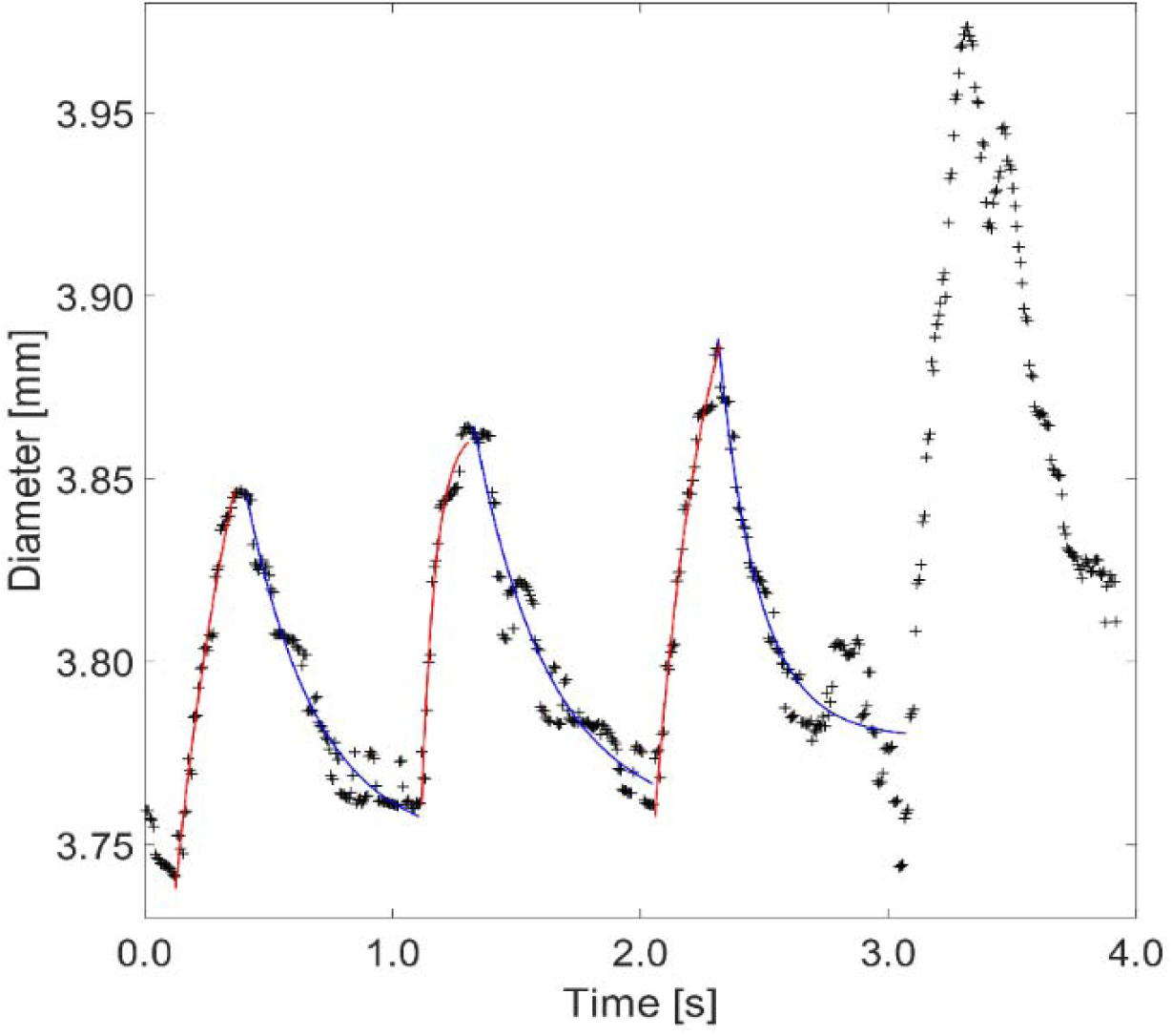
Diameter variation as a function of time at a specific cross section showing four cardiac cycles. The crosses show the measured results and the lines are the fitting curves for dilatation (red) and relaxation (blue). The diameter of the last cardiac cycle in the figure was not well fitted by a two time-scale decay and these data sets were excluded and not included in further analysis.

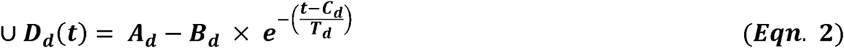

Where ***A*** and ***B, C*** and ***T*** are parameters to be fit. The relaxation during diastole was not well modelled by one time constant and therefore two time constants were employed to fit the diameter.

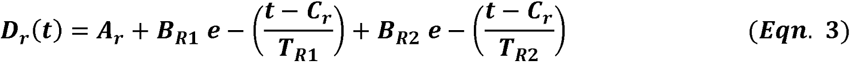

Here the relaxation time constants will be referred to as fast, T_R1_ and slow, T_R2_. The motivation for using exponential modes is that tissue is commonly modelled using as a series of springs and dashpots, the so-called Zener model which has exponential solutions ^11^. For systole the response is a convolution of the driving pressure and vessel wall dynamics and so extracting wall properties is challenging. However, for diastole the temporal dynamics should be dominated by the properties of the wall which are captured by T_R1_ and T_R2_. The slow fall occurs under minimal arterial pressure and so is most sensitive to wall properties.

Assuming the incoming pulses are square waves with very short on duration, the tissue response is formed by three compartments in terms of the diameter: (i) steep rise (a convolution of the driving waveform and the tissue response), (ii) steep fall, (iii) slow fall before the next pulse comes. The steep rise and fall are caused by the forcing input, thus the composition of these two parts are mainly the input forcing term.

The best fit curves and time constants were constructed using the `fminseaTch` function in MATLAB® (The MathWorks, Inc.), based on the Nelder-Mead simplex algorithm ^12^, in which convergence in low dimension has been verified in ^13^. The cost function that is used in the minimalisation is the summation of standard L2-Norm, between the data and either Equation 2 or Equation 3.

The initialisation of parameters for `fminsearch` was as follows: The vertical (A_d, A_r) and horizontal (C_d, C_r) shifts’ were set to the mean of the values’ range within a fitting section to start with (diameter and time, respectively). The exponentials’ magnitudes (B_d, B_r and D_r) were the size of one pixel. *T*_*R*2_ was set to the time period of the relaxation section (ie diastole) and *T*_*R*l_ *T*_*R*2_ / 10.

The brachial artery relaxation index (BARI) is defined by the mean relaxation constant of the exponential curves. The pilot data indicated that *T*_*R*2_ was more likely to be a parameter differentiating aneurysm growth rate. Therefore, resulting analysis was focused on the diameter relaxation sections and the corresponding fittings.

## Results

Algorithm defined lumen wall positions were faithful to true brachial artery wall position in good quality ultrasound recordings (**Figure 2**, panel A) with an accuracy on the order of an image voxel (21 µm). The algorithm was unable to reliably locate wall position for 14 patients and these were excluded from further analysis. The most common issue encountered was the presence of a slice artefact in the lumen (**Figure 2**, panel B).

Matlab was used to create a 3D plot (**Figure 3**) displaying vessel wall movement, as a function of position along vessel and time through recording. As the segmentation algorithm decides how complicated the contour’s shape must be to represent it at certain accuracy, the number of points representing each contour is generally different for a pair of top and bottom contours. In some cases, the segmentation returns “S” and “Z” shaped contours, means that for a single (x) position entry, there may be more than one contour (y) position values. The algorithm was written to include feasible manipulations to mitigate the above two issues in Equation 1. Before calculating the diameter, each contour is checked for whether there is more than one entry on a single horizontal position. If there is, the average is taken to be the value used in Equation 1.

For every location along the artery the diameter change was captured during cardiac cycles and curves fitted for dilatation and relaxation (**Figure 4**).The generated time constants were hypothesised to distinguish between fast and slow growing AAAs.

A pilot study was performed on six patients with the slowest (S1-3) and fastest (F1-3) growing aneurysms in the OxAAA cohort. The time constant for diameter dilatation sections (*T*_*D*_)was not different between the two groups. BARI values are shown as bars in **Figure 5** (green for slow growth AAA patients, red for fast growth AAA patients). The data suggests that BARI can be used to separate fast from slow based on a BARI threshold of 1.5s.

**Figure 5.**
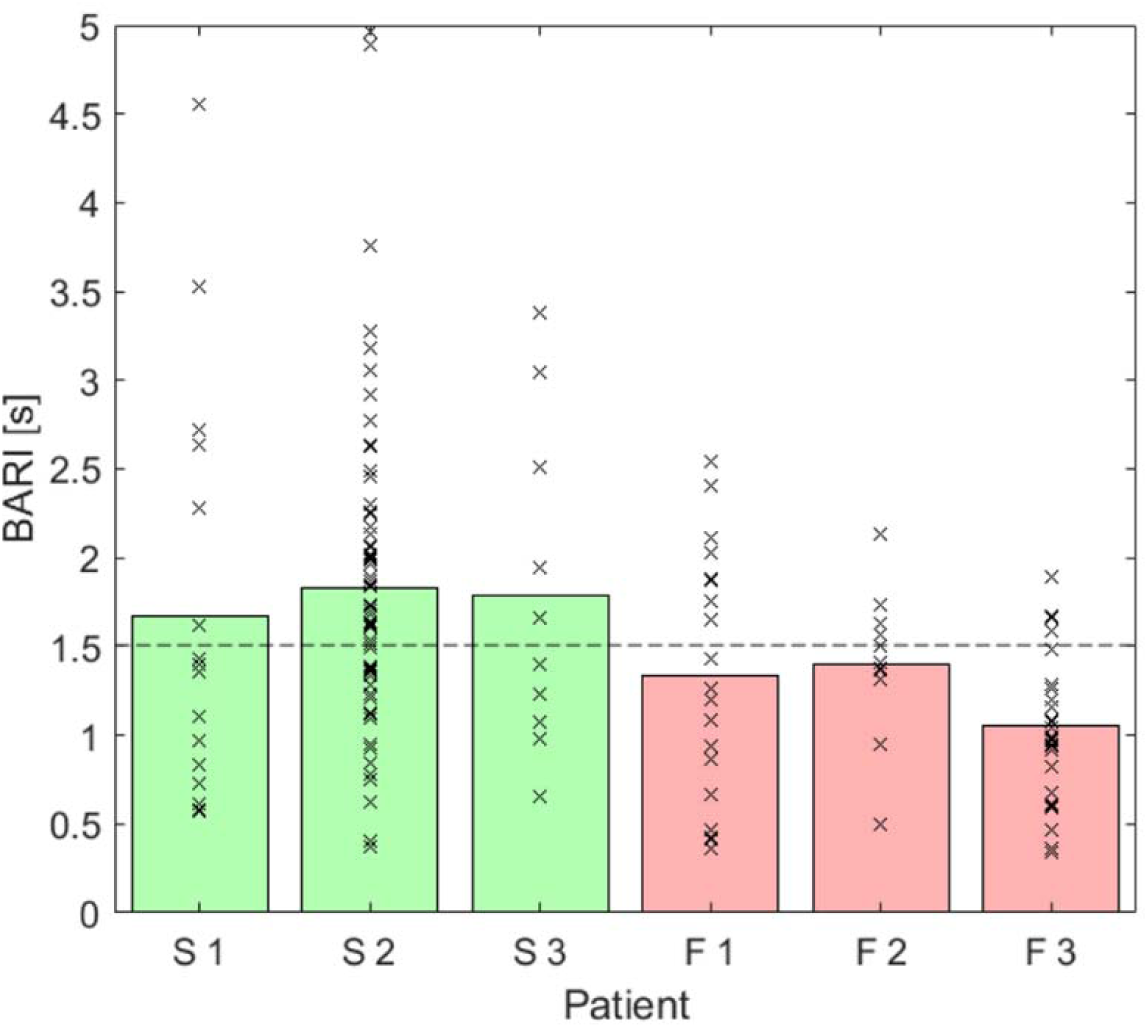
Comparison of BARI between representative slow (S1,S2 and S3) growth and fast growth (F1, F2 and F3). Relaxation time constant values are marked in crosses for each patient. The bars represent the BARI values derived from the data crosses for each patient (Green: slow growth patients. Red: fast growth patients). Dashed line at BARI = 1.5 suggests a threshold value to separate patient’s growth type.

This positive result demonstrating separation in a pilot study for the patients at the extreme spectrum of prospective AAA growth and the plausible theoretical basis for the biomarker led to further analysis being performed on rest of the OxAAA cohort.

### Difference between BARI in healthy volunteers and aneurysmal patient cohort

16 healthy volunteers were age and gender matched to 16 patients from the aneurysmal cohort to assess whether BARI differed between patients with AAA (cohort) and without aneurysms (HV). There was no statistical difference observed between the two groups on unpaired t-test (HV median 0.93, HV interquartile range 0.65-0.99, Cohort median = 0.89, Cohort interquartile range = 0.82-1.03, P=0.78).

### Difference between BARI in aneurysms of different sizes

Patients from the OxAAA cohort (n=124) were divided into three subgroups according to their aneurysm AP diameter: small (30-39mm), moderate (40-55mm), and large (>55mm). There was no significant difference observed between groups on Kruskal-Wallis (Small vs Large, P=0.82, Small vs Medium, P=0.99, Medium vs Large, P=0.99). Spearman correlation of BARI against aneurysm size (mm) was not significant (p=0.11), **Figure 6**.

**Figure 6.**
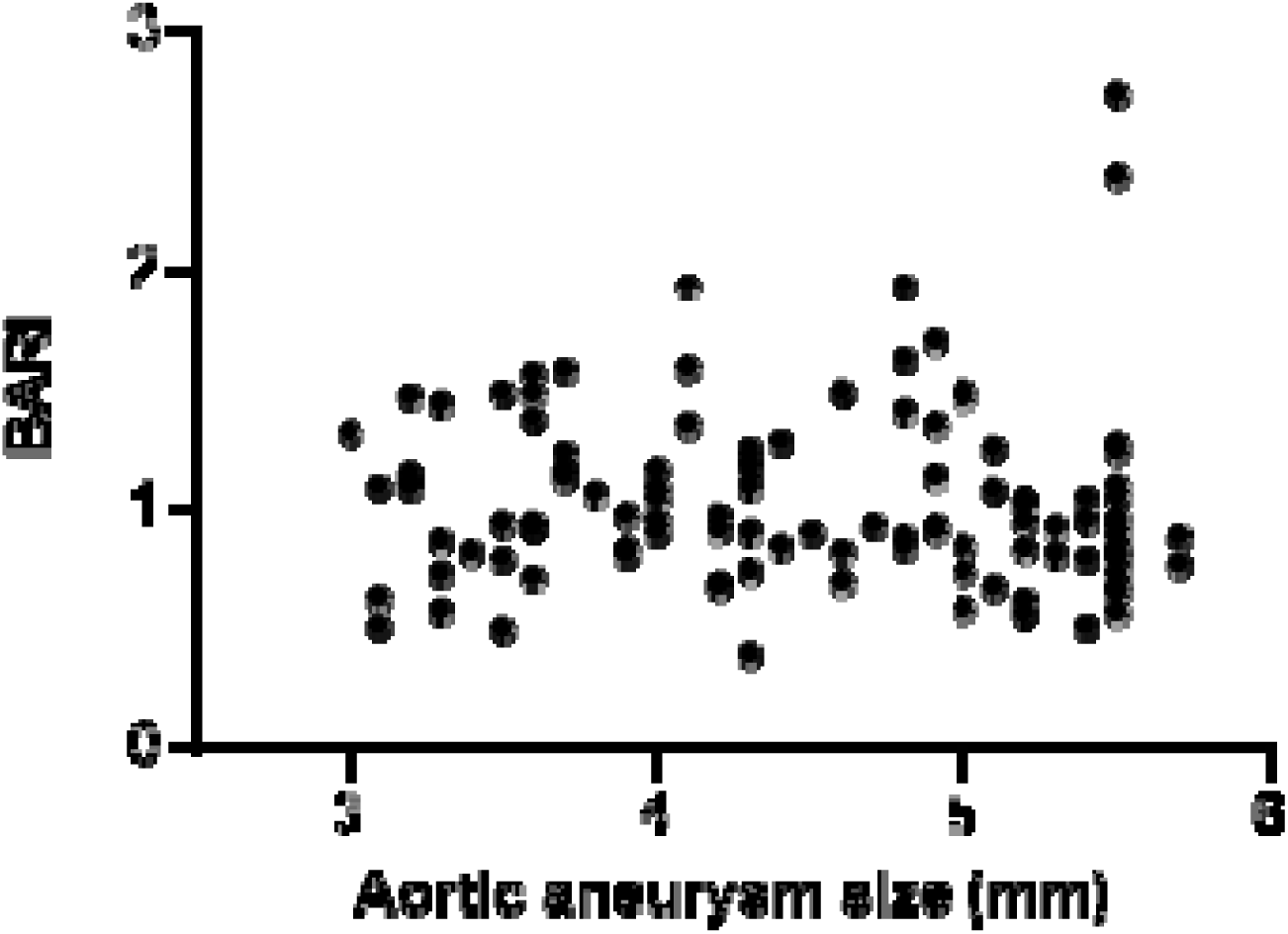
BARI vs AAA size for 124 patients. No correlation between aneurysm AP diameter (x-axis) and BARI (y-axis) P=0.68

### Correlation between BARI and future AAA growth

The growth rate of all subjects in the OxAAA cohort (n=124) was evaluated over 12 and 24 months dependent on duration of size data collected for each subject. **Figure 7** shows measured BARI for each patient versus the AAA growth and no correlation was. Spearman correlation of BARI against % aneurysm growth per year was not significant (p=0.45).

**Figure 7.**
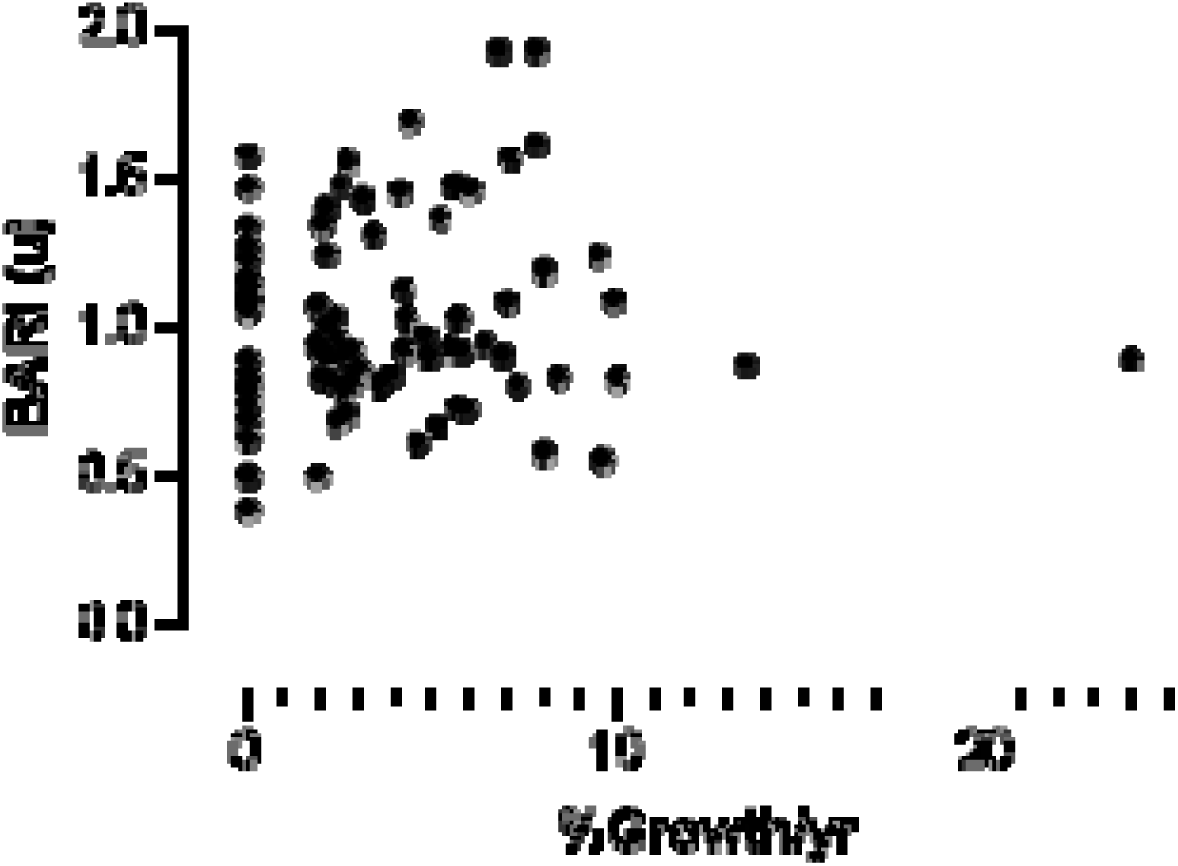
BARI vs AAA % growth per year for 124 patients. Linear regression comparing aneurysm % AP diameter growth per year (x-axis) and BARI (y-axis) Slope 0.01 P=0.46

## Discussion

This work was motivated by the desire to predict the growth and progression of AAAs. The existing surgical threshold based on a size criteria (>55mm) has intrinsic shortcomings, as aneurysm size is not an absolute predictor of the risk of rupture, and the rate of AAA progression varies greatly between individuals. Those with moderate size aneurysms have a small (1%) annual risk of rupture and the majority of patients (70%) with an initially small or moderate size AAA will progress and require surgery within 5 years ^4, 5^. Improved biomarkers of future AAA progression would enable the stratification of patients who may benefit from surgery earlier than the existing size threshold and decrease the perioperative risk to those patients.

Our hypothesis was that US based measurements of the deformation of vessels walls could be used to extract mechanical properties of the vessels to provide information that could improve the ability to predict AAA progression. Preliminary data suggested that BARI, a measure of a relaxation time scale had the most promise. Unfortunately, there was no significant difference in BARI between aneurysmal patients and healthy volunteers, nor did it vary according to aneurysm size. It offered no predictive value for aneurysm growth at 12 or 24 months. Although BARI did not appear to improve the ability to predict growth in AAAs, the processing approach described here to extract mechanical properties from ultrasound image analysis may have alternate applications in other pathologies or the research setting.

Investigating physiological characteristics of vasculature using imaging is an important part of developing our understanding of cardiovascular and aneurysmal disease. Better definition of the biophysical characteristics of vasculature in healthy and pathological states is important to help underpin future research. Ultrasound is an ideal imaging modality to extract biomechanical properties as it is non-invasive, functional, widely accessible and without exposure to ionising radiation.

We had hypothesised BARI would help the predictive index for aneurysm growth, in addition to existing biomarkers such as FMD, AAA diameter and circulating plasma proteins ^9, 14^. Our data did not support this. It would have been preferable to perform the same analysis using ultrasound imaging of the AAA itself. However, as the AAA is an organ structure deep inside the abdominal cavity, ultrasound imaging of the AAA is only feasible using a low frequency probe (typically at 3MHz) to achieve the depth penetration. The images acquired using this probe were found to be too low in resolution for the analysis described here to extract pulsatile expansion of the AAA. Nevertheless, BARI is a novel image analysis technique which underscores the importance of active pursuit of imaging biomarkers, across various modalities to help inform clinical decision making for AAAs.

## Acknowledgement

The Oxford Abdominal Aortic Aneurysm Study was supported by the following: University of Oxford, Medical Sciences Division Medical Research Fund (MRF/HT2016/2191); University of Oxford, Nuffield Department of Surgical Sciences; John Fell Oxford University Press Research Fund (142/075); National Institute of Health Research (NIHR) Oxford Biomedical Research Centre; RL was supported by a Academy of Medical Science Starter Grant, UK (SGL013/1015). PL was supported by an EU Erasmus+ traineeship studentship. AC is a Clarendon Keble Scholar, University of Oxford. Work described in this manuscript is subject to a international patent filing (WO/2020/030890).

Contributors to the OxAAA Study include: Amy Jones, Felicity Woodgate, Nicholas Killough, Kirthi Bellamkonda, Sowmya Mangipudi, Members of the Oxford Regional Vascular Services and the Jackie Walton Vascular Studies Unit.

